# Genomically integrated orthogonal translation in *Escherichia coli*, a new synthetic auxotrophic chassis with altered genetic code, genetic firewall, and enhanced protein expression

**DOI:** 10.1101/2023.11.18.567690

**Authors:** Hamid Reza Karbalaei-Heidari, Nediljko Budisa

**Affiliations:** Department of Chemistry, University of Manitoba, Winnipeg, MB, Canada; Institute of Chemistry, Technical University of Berlin, Berlin, Germany

**Keywords:** CRISPR-associated transposase tool (CASTs), genetic code expansion, genomic integration, genetic firewall, synthetic auxotrophy, orthogonal translation systems (OTS)

## Abstract

In the last three decades, genetic code engineering has expanded protein biosynthesis options from the natural set of 20 canonical amino acids to over 250 non-canonical amino acids (ncAAs). This progress involves rewiring of protein translation by establishing Orthogonal Translation Systems (OTS) through orthogonal pairs. Traditionally encoded on plasmid vectors, these systems are often unstable and burdensome in large-scale fermentations. To moving forward from academia to reliable technology, it is crucial to integrate OTS genetic modules stably into a molecular chassis with a defined genome background. Here, we demonstrate genomically integrated OTS in *Escherichia coli*, creating a synthetic auxotrophic chassis with an altered genetic code. Using CRISPR-associated transposase tool (CASTs), we targeted multiple genome sites, inserting OTS components (enzymes, tRNA genes) non-disruptively. Our OTS system demonstrated site-specific incorporation of *m*-oNB-Dopa through in-frame amber stop codon readthrough, enabling the expression of smart underwater bioglues. Simple metabolic labelling, introducing fluoroproline analogs enhancing conformational stability during orthogonal translation, further bolstered system robustness. These chassis, equipped also with synthetic auxotrophy for *m*-oNB-Dopa, serve as a built-in genetic barrier (genetic firewall), ensuring safe bioproduction in genetically isolated settings.

**Graphical Abstract:** 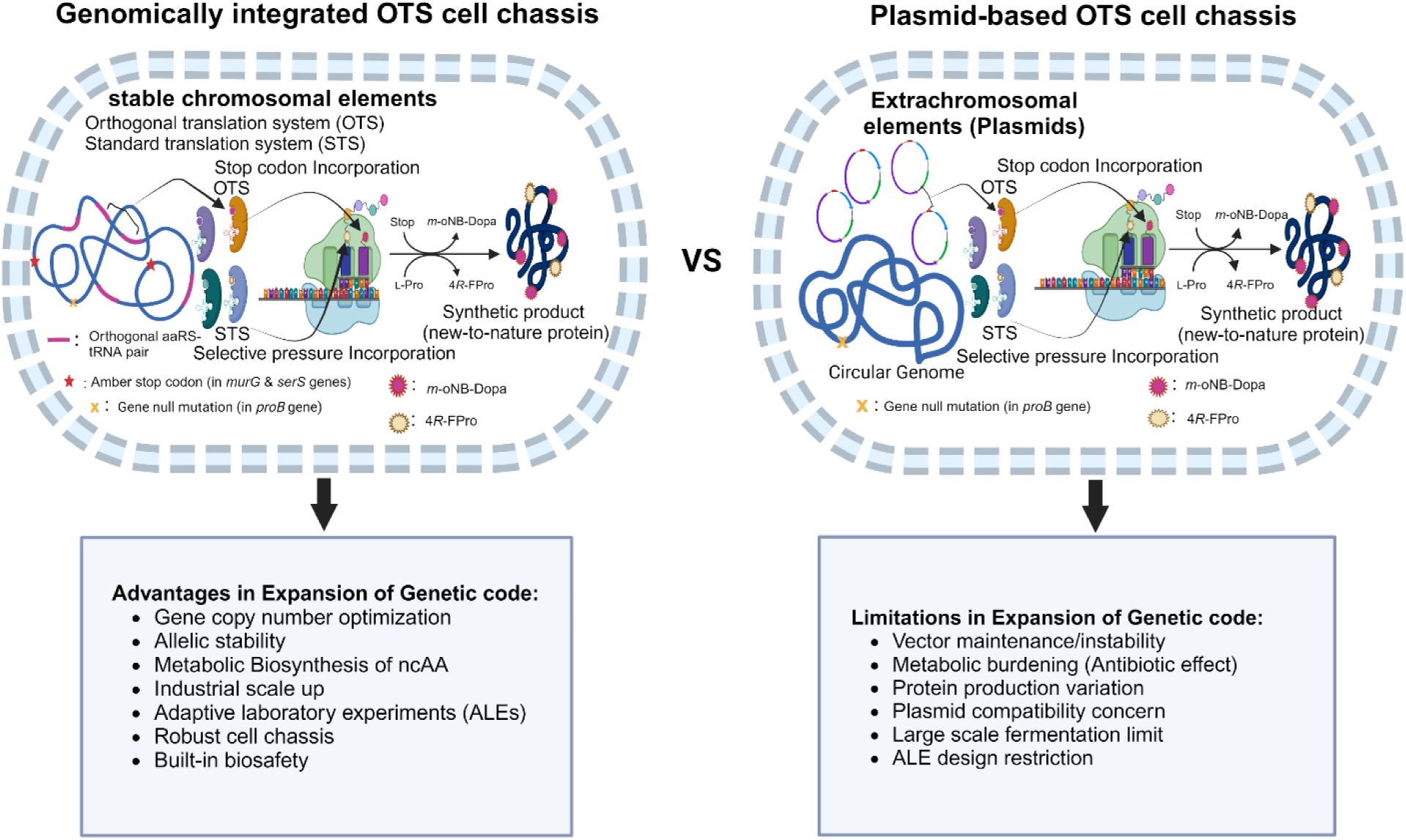

**One-sentence Abstract:** CRISPR-assisted transposition has enabled the development of a robust and biosafe E*scherichia coli*-based chassis with genomically integrated orthogonal translation components, enhancing synthetic protein production and laying the foundation for the transition of this research field from academia to reliable technology.

## Introduction

Nature uses a limited number of building blocks for nucleic acids and proteins in biology. While there are four nucleotides for nucleic acids and 20 canonical amino acids (cAAs) for proteins, the reprogramming of ribosomal translation allows the incorporation of non-canonical amino acids (ncAAs) which confers different chemical and biophysical properties to proteins. This technology has immense potential as it enables the precise placement of different ncAAs in proteins *in vivo* and can be used in new drug development, bio-orthogonal engineering, customisation with specific molecules and the introduction of post-translational modifications to study cellular regulatory processes or even to create synthetic cells with unique genetic codes^1^. To establish reliable chassis as robust platforms for this technology, it is important that they are genetically distinct from their natural ancestors. Improving their resilience can be achieved through genomic recordings^2^ and methods such as Adaptive Laboratory Evolution^3^. These synthetic cells are equipped with a genetic firewall and strict biocontainment measures to reduce genetic exchange with the natural environment and ensure build-in biosafety in biotechnological applications ^4^.

Toward these goals, gene libraries of selected aminoacyl-tRNA synthetases (aaRS) are being created for orthogonal translation systems (OTSs). Natural archaeal systems such as the *Methanocaldococcus jannaschii* tyrosine pair (*Mj*TyrRS:tRNA^Tyr^) and the *Methanosarcina mazei or bakeri* pyrrolysine pair (*Mm/b*PylRS:tRNA^Pyl^) are rationally engineered and used to select new OTSs in *Escherichia coli* (*E. coli*)^5^. These orthogonal aaRSs, together with compatible suppressor tRNAs and other engineered translational components, are usually encoded on plasmid vectors. However, a major challenge with these systems is their instability when culturing bacteria on a large scale. Plasmid-free cells tend to proliferate and overgrown as the host cell population grows, which is problematic for biotechnological applications^6^. Ensuring a consistent gene dosage is essential for effective protein synthesis and the sustained resilience of synthetic cells across various environments.

Hence, there is a significant need for the stable and secure integration of OTS components into the host genome^7^. This necessitates an efficient chromosomal integration method for precise localization, controlled gene dosage, and genetic stability. Ideally, a safe and genetically stable cell-based synthetic chassis equipped with OTS should possess genomes that integrate orthogonal translation components. This integration enables the chassis to operate within an expanded genetic code, incorporating ncAAs and leveraging their unique metabolism for in-site biosynthesis ^8^.

Genome editing tools enable the simultaneous and safe insertion of orthogonal pairs and other OTS-related genes as well as selected metabolic markers into the genome of a chosen cell chassis. An example of this capability would be the integration of an orthogonal translation system (OTS) and desired metabolic markers in *E. coli*, a widely used laboratory microorganism. This enables *E. coli* to synthesise the desired ncAAs internally and to produce synthetic proteins from simple metabolic precursors or substrate analogues^9^. To date, several strategies for integrating gene cassettes in *E. coli* have been described, each with its advantages and disadvantages^10,11^. One of the main obstacles in establishing integrated OTSs is the difficulty in accurately locating them in the genome and adjusting their gene dosage or expression levels. In general, reported OTSs are less efficient than the components of the native translational machinery^12^, and despite endogenous aaRSs, which generally have high catalytic efficiency, we also need to regulate and target gene expression (among others) at the gene copy number level.

Recently discovered CRISPR-associated transposons offer a promising approach for achieving a genetic firewall in synthetic cells. These transposons, such as Type I-F, I-B, and V-K CRISPR-Cas systems, have adopted an RNA-guided DNA insertion strategy that doesn’t create double-strand breaks (DSBs) ^13,14^ which rely on direct homology repair mechanism. Three terms in the literature describe this technology: CRISPR-ASsisted Transposition (CAST)^15^, INsertion of Transposable Elements by Guide RNA-Assisted Targeting (INTEGRATE) ^16^, and MUlticopy Chromosomal Integration by CRISPR-Associated Transposases (MUCICAT) ^17^. CAST is a subtype V-K of Tn7-like transposons found in *Scytonema hofmanni*. It uses the Cas12K CRISPR-Cas effector but has some limitations, including low integration efficiency and frequent insertion of the entire donor plasmid due to the lack of the transposase A (TnsA) enzyme ^18^.The INTEGRATE and MUCICAT systems are derived from a type I-F CRISPR-associated Tn6677 transposon from *Vibrio cholerae*. They contain TnsABC, TniQ, and a CRISPR-Cas multi-protein complex cascade composed of Cas8, Cas7, and Cas6. These systems offer high integration efficiency with minimal detectable off-target effects or other integration byproducts^16^.

By utilizing a CRISPR array with consecutive spacers directed at various positions within the target genome and flanked by an upstream 5’-CC-3’ PAM sequence, it’s possible to simultaneously target multiple loci on the chromosome^19^. These targets allow the insertion of a desired DNA fragment (Cargo) approximately 49 bp downstream of the target sequence. Each type I-F CRISPR array comprises two specific repeats (28 bp) and a unique spacer (32 bp). Targeting multiple sites in the genome is crucial for efficiently integrating the OTS. The CAST system’s advantage lies on its ability to insert the desired DNA fragment without causing double-strand breaks in the chromosome, which makes it less disruptive to existing coding sequences. Following transposition experiments, screening integrated strains with various copy numbers of the cargo genes provides ample opportunities to optimize the functionality of the desired OTS.

We harnessed the Type I-F Tn7-like transposase from *Vibrio cholera* (Tn6677) as a highly efficient tool to integrate the OTS into recoded *E. coli* B-95.ΔA, specifically the B-95.ΔA NK53 strain^20^, which is a nitroreductase-deficient strain derived from the *Escherichia coli* B95ΔA strain (originally derived from *E. coli* BL21(DE3))^21,22^. In this study, we used the integration of an orthogonal *Mj*TyRS (oNB-DopaRS)-tRNA_CUA_ cassette (O-pair) as an example of OTS integration in *E. coli* for site-specific incorporation of *m*-oNB-Dopa (*meta*-(ortho-(2-nitrobenzyl))-3,4-dihydroxyphenylalanine)^23^ by in-frame amber stop codon readthrough. We optimized the copy number of a gene cassette (pTacI-oNB-DopaRS-*proK*-tRNA_CUA_) within the engineered *E. coli* genome. This optimization was achieved using an array of crRNAs with nine target spacers. To assess the efficiency of our engineered cell chassis, we tested it by expressing Sumo-sfGFP, which contains one or three TAG amber codons, and elastin-like protein (ELP) constructs with varying numbers of amber codons (5, 10, 20, and 30 TAGs) fused to superfolder Green Fluorescent Protein (sfGFP) at different scale of cultures.

**Scheme 1.**
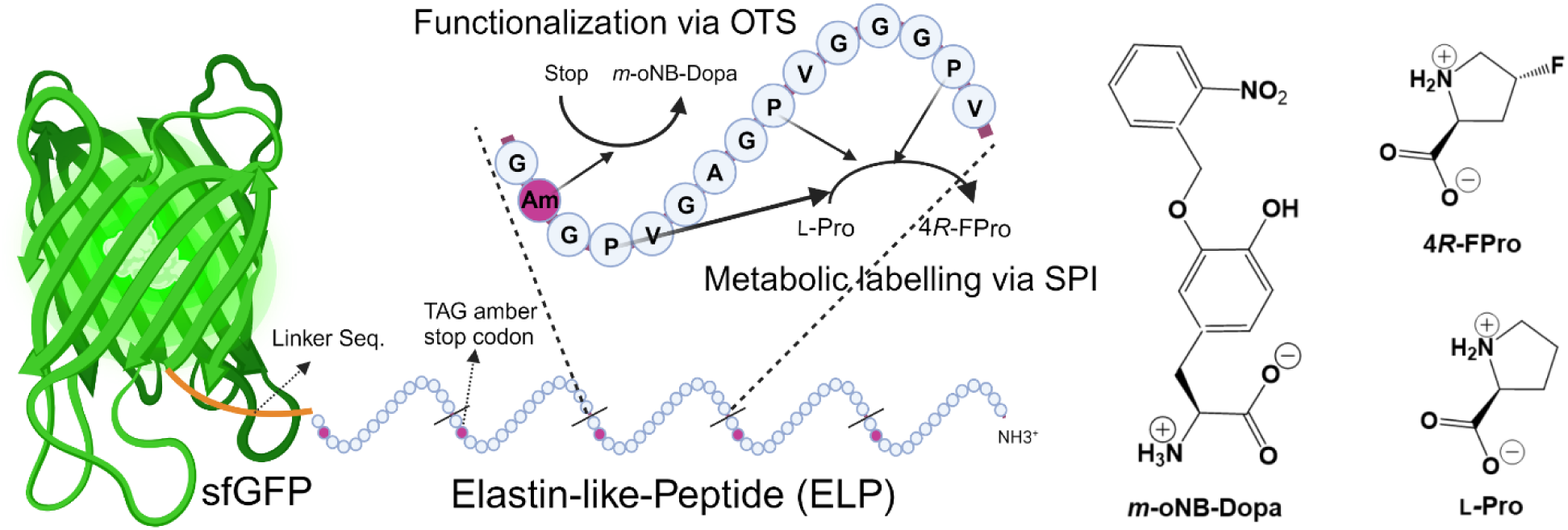
ELP(5xAm)-sfGFP model protein was expressed in BNH24 (Δ*proB*), MS34 strain, in autoinduction minimal medium containing 1 mM 4R-FPro and 2 mM *m*-oNB-Dopa. Created with BioRender.Com.

We have also advanced an OTS-integrated strain by knocking out the *proB* gene, creating a proline auxotroph as a metabolic prototype that can further expand the scope of reprogrammed protein translation. This allowed us to incorporate non-canonical proline-like amino acid analogues using a method called selective pressure incorporation (SPI)^24^ in parallel to orthogonal translation. We specifically targeted the incorporation of three proline analogs, 4*S*-fluoroproline (4*S*-FPro), 4*R*-fluoroproline (4*R*-FPro), and 4,4-Difluoroproline (Di-FPro), which influence protein backbone conformation and translation rate^25^ to see their effects on orthogonal translation. 4*S*-FPro promotes peptide bond formation in *cis* conformations and leads to ribosome stalling and a reduction in protein yield, while 4*R*-FPro favour *trans* conformations, accelerating translation and consequently leading to increased expression of the target protein^26^. To this end, we employed a proline auxotroph chassis to integrate residue-specific sense codon reassignments, substituting Pro residues with (4*S*/*R*)-FPro analogues, along with the position-specific insertion of *m*-oNB-Dopa at in-frame amber stop codons in ELP (*n* × TAG)-sfGFP (*n* = 5, 10, 20, and 30) via translational readthrough. All of this was achieved in a single *in vivo* expression experiment (Scheme 1).

Finally, we established a genetic firewall in these biocontained OTS cells. This entailed introducing amber (TAG) codons into the conserved Phe residues of essential genes, *murG* and *serS*^4^, while ensuring robust growth in a permissive medium (LB+*m*-oNB-Dopa). These measures resulted in synthetic auxotrophy with an undetectable escape frequency when culturing around 10^9^ cells in LB media, both solid and liquid, ensuring the full biosafety of our chassis.

## Results

### Generation of an efficient integrated cell chassis based on INTEGRATE and MUCICAT

To improve efficiency of the multiple integration approach, we tried to change the copy number of the plasmids participating in the transposition event. Original technique was based on three-plasmid system, whereby pQCascade-array1 encoded *Vch*TniQ, *Vch*Cas8, *Vch*Cas7, *Vch*Cas6, and crRNA-array1, and the pSL0283 plasmid encoded TnsA, TnsB, and TnsC, where the expression of all RNA-protein components was under the control of T7-lacO promoter using a medium copy number (CloDF13 ori), and low copy number (ColA ori) origin of replication, respectively. The third donor plasmid had a high copy number ori (ColE1) containing 127 bp and 134 bp right and left transposon ends, respectively ^9^. In an improvement study, Zhang et. al. replaced T7-lacO promoters with the Ptet promoter. The Ptet promoter is a tight inducible system that segregates the regulation of the elements of the transposition machinery from the cargo genes, induced by IPTG ^12^. Nonetheless, the existing system encounters issues with the transformation of the pDonor plasmid, resulting in slow growth of transformants and an overgrowth of unproductive mutants on the transposition plate. To address these challenges and enhance transposition efficiency, Vo and co-workers suggested using of a single pSPIN construct incorporating the pBBR1 vector backbone. This new approach exhibits an integration efficiency of over 90% for a single spacer and a stronger preference for insertion events where the transposon’s right end is closer to the target site^11^. However, single plasmid system still suffers from difficulties in cloning of large cargo genes and limits the manipulation of mini Tns.

Our approach involves a two-plasmid system. The first plasmid, using the CloDF13 ori as a medium copy number vector backbone, employs the tetA/tetR promoter for tightly regulated induction of transposition components (including the crRNA array and the polycistronic messenger RNA for the multiprotein complex) while the second plasmid, with a ColA ori vector backbone, carries the cargo genes in the form of mini-Tn (see Fig. 1). The expression of the coding sequences of pTet-Effector-array1 (PM452) is controlled by the concentration of anhydrotetracycline (aTc) and targets nine distinct genomic sites through the transcription of a multispacer CRISPR array.

**Fig. 1.**
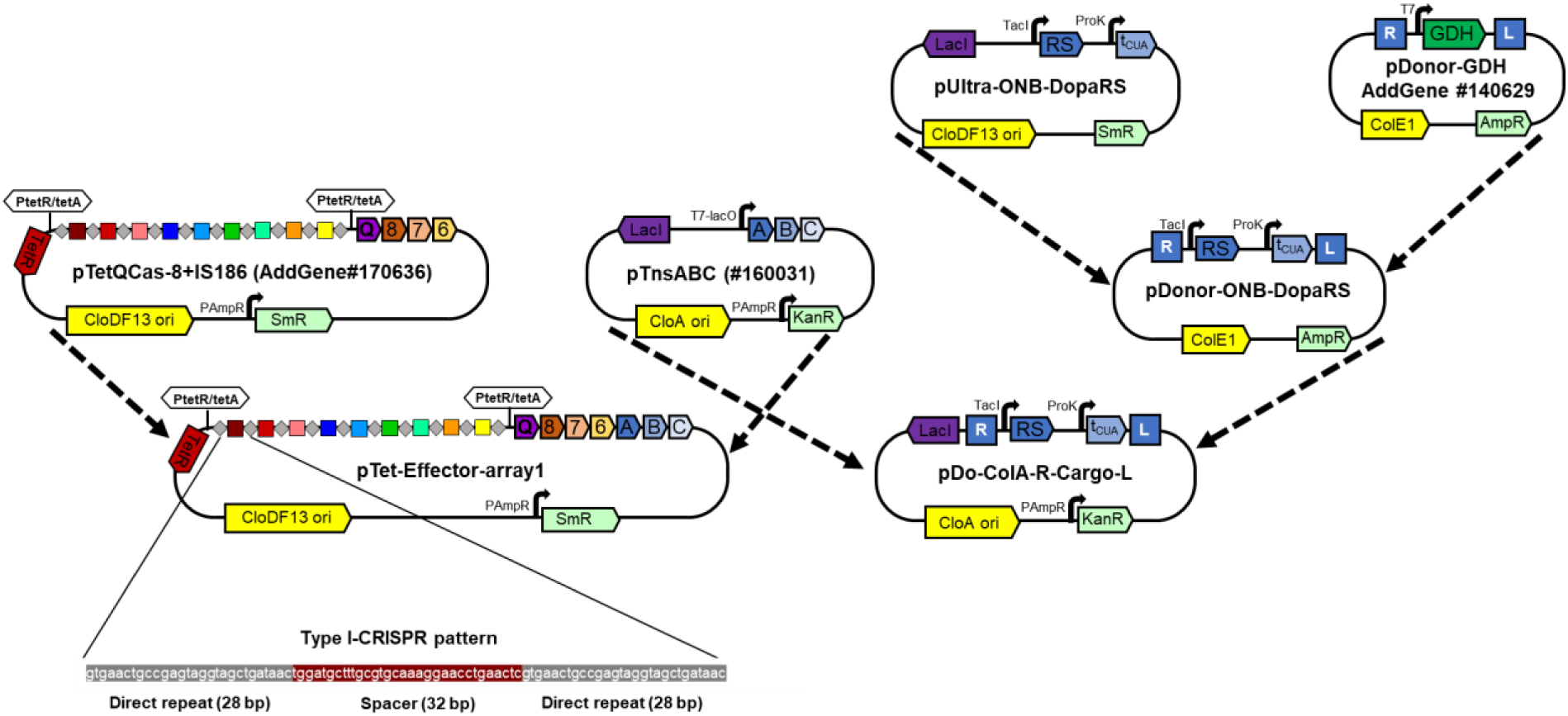
Schematic representation of two plasmid-based transposition vector constructs. The pDo-ColA-R-Cargo-L vector was generated through Gibson assembly, combining DNA fragments from the pTnsABC (Addgene#160031) vector backbone and the pDonor-oNB-DopaRS mini-Tn region. Initially, the pDonor-oNB-DopaRS vector was derived from the pUltra-oNB-DopaRS and pDonor-GDH vectors (Addgene#140629). To construct the pTet-Effector-array1 vector, the amplified polycistronic TnsABC gene fragment from the pTnsABC vector (#160031) was cloned in the pTetQCas-8+IS186 vector (Addgene#170636), using Gibson DNA assembly technique. The pattern of type I-CRISPR crRNA sequence for the IS4-like element was expanded to emphasize the architecture of the array1 crRNAs.

The pTacI-oNB-DopaRS-proK-tRNA_CUA_ gene cassette, referred to as OTS, was inserted into the pDo-ColA-RE-Cargo-LE vector, creating a new construct named pDo-ColA-oNB-DopaRS-tRNA_CUA_ (PM453). When this double-plasmid transposition was performed in an engineered *E. coli* strain, NK53 (derived from B-95.ΔAΔfabR), it yielded a satisfactory number of colonies after an overnight incubation at 37°C. The results validate that the newly engineered transposition plasmids exhibit a reduced negative effect on both bacterial growth and the transposition event. Inducing transposition by adding 1-5.0 ng/mL of anhydrotetracycline (aTc) resulted in a range of insertion events following overnight incubation at 37°C, varying from single to triple gene cassette integrations in the first round of transposition, as detected by PCR (see Fig. S1). Subsequent plasmid removal/curing led to the identification of strains, which were named based on the number of OTS insertions. This naming convention started with strain BNH21 for a single insertion and went up to strain BNH23 for triple OTS cassette insertions.

### A cell chassis with OTS integration for effective non-canonical amino acid incorporation

To assess the efficiency of incorporating the ncAA *m*-oNB-Dopa into the OTS-integrated strains, we used fluorescence measurements on the Sumo-sfGFP model protein, which harbored either one or three Amber (TAG) codons. We employed fluorescence intensity as an indicator of protein production levels. When examining a protein with a single Amber codon, it exhibited a substantial background signal due to natural suppression, making it challenging to differentiate between the efficiency of various strains (see Fig. 2). However, when we analyzed overexpression of a protein containing three Amber codons, we observed a direct relationship between the number of OTS integrations in the strain’s genome and an increase in fluorescence signal (see Fig. 2).

**Fig. 2.**
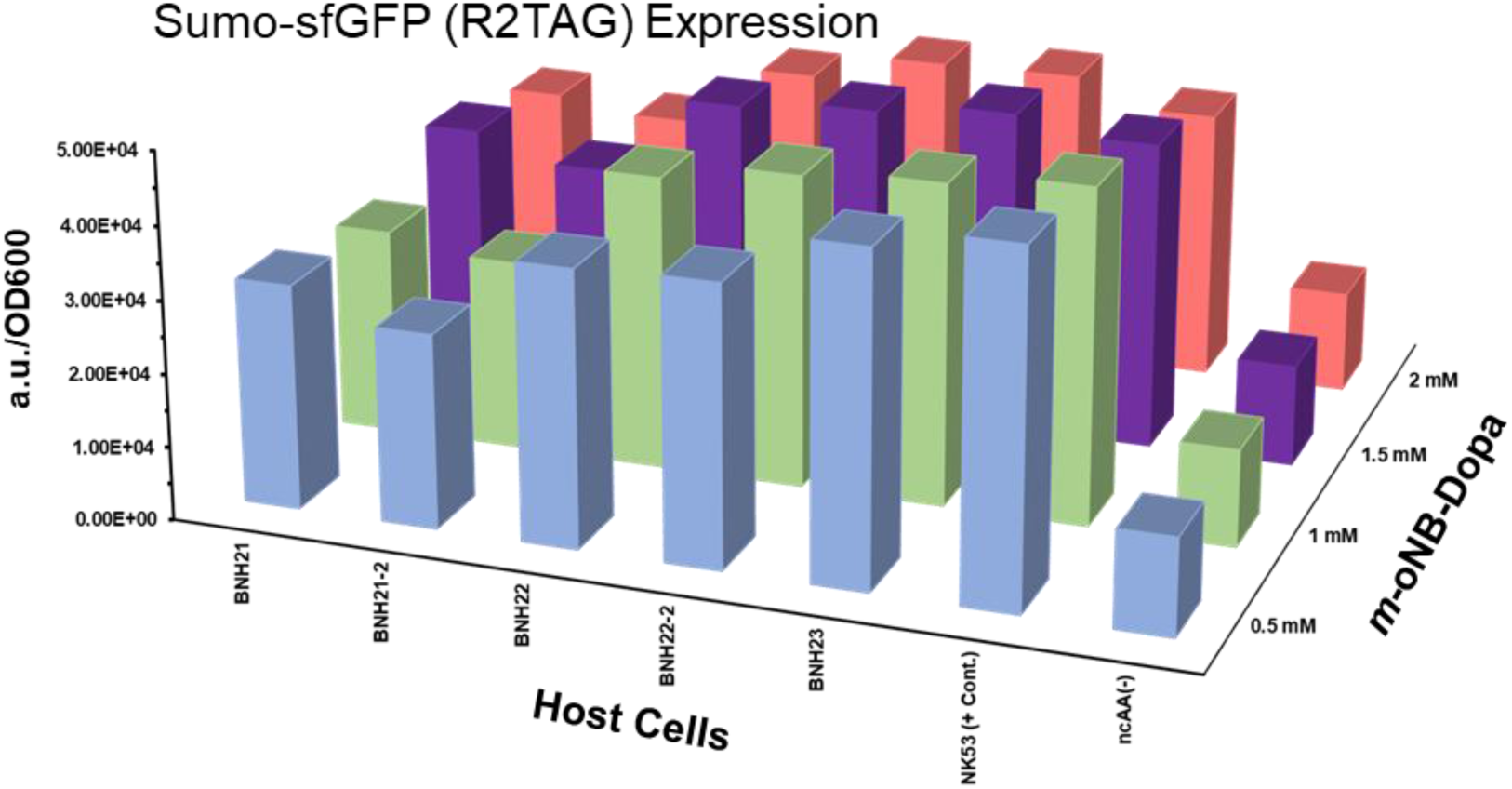
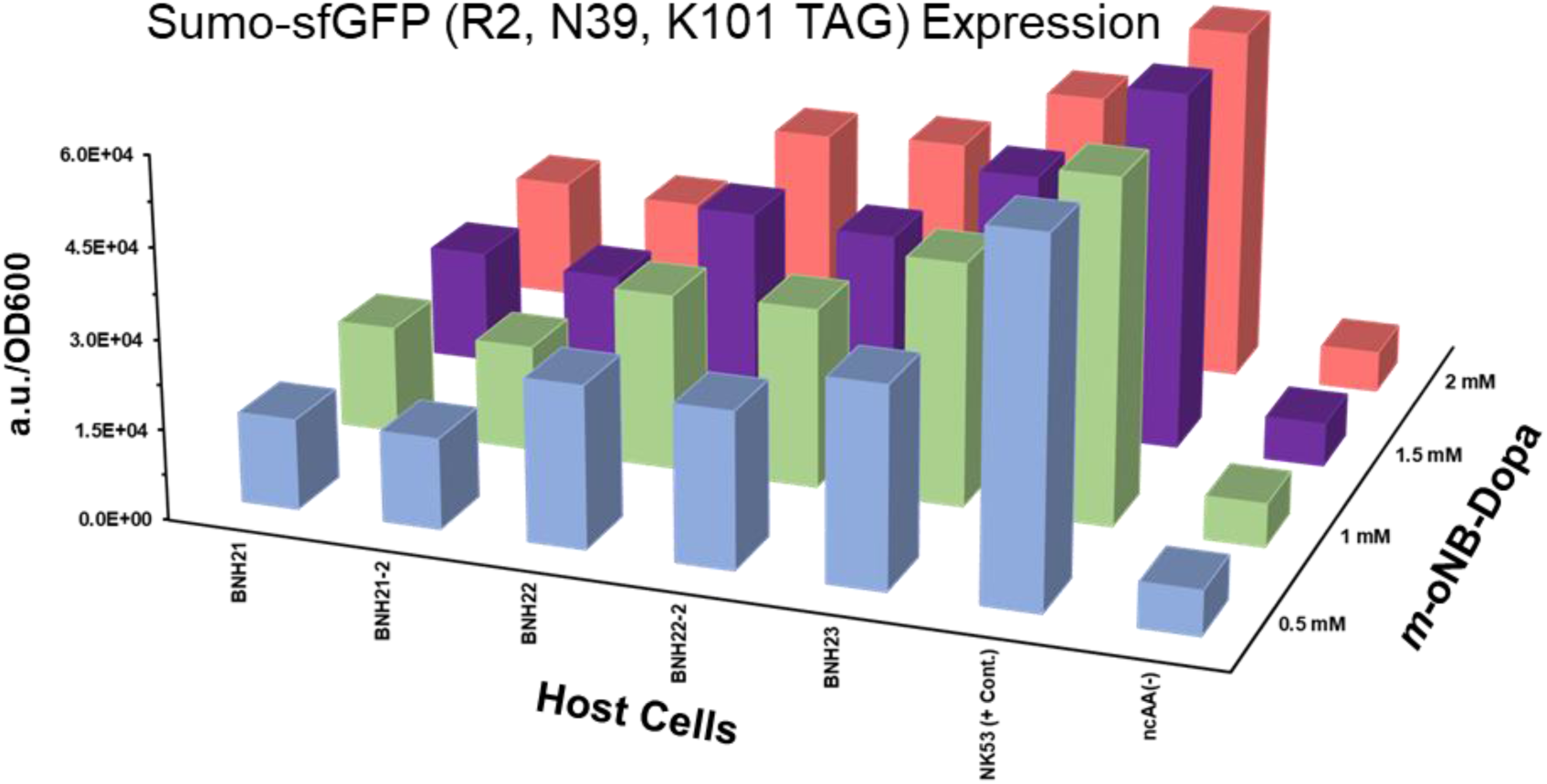
Analysis of the microplate fluorescence signal as a marker for the efficiency of *m*-o-NB-Dopa incorporation. Strains harboring pET28b-Sumo-sfGFP plasmids with either R2TAG or a combination of R2, N39, K101TAG were cultivated in 100 μL ZYP-5052 medium with varying concentrations of *m*-oNB-Dopa analog (0.5-2.0 mM). The culture was incubated in a 96-well microplate overnight at 37°C and 300 rpm, and the fluorescence intensity per OD_600_ (indicating bacterial growth) was determined as a calibrated signal. The microplate was excited at 488 nm, and the fluorescence signal was measured at 530 nm using the Tecan Infinite 200 PRO plate reader. Error bars not shown. Data have been presented in Supplementary information as mean ± SD.

Fluorescence intensity measurement of the cultures exhibited a linear increase from BNH21 to BNH23, corresponding to single to triple integrations of the oNB-DopaRS:tRNA_CUA_ o-pair cassette. This effect was particularly pronounced when three Amber codons were introduced into Sumo-sfGFP (R2, N39, K101Am). We also conducted experiments to assess the impact of the integration site in the genome on incorporation efficiency, comparing two strains with one copy and two strains with two copies of the OTS (BNH21, and BNH21-2 or BNH22, and BNH22-2) at different genome sites. The results confirmed that there were no significant functional differences in the integrated o-pair at various target sites.

### Transposition enrichment

The enrichment of cassette integration was achieved through repeated transposition events in strain BNH23. During the second round of transposition, we isolated two strains that had integrated four and five copies of the cargo gene cassette, which we designated BNH24 and BNH25, respectively (Fig. S2). When we compared the incorporation of *m*-oNB dopa between the modified chassis and the strain NK53, which had the double plasmids pUltra-ONB-DopaRS-tRNA_CUA_ and pET28b-Sumo-sfGFP (3xAm), we observed that after integrating five copies of the cassette into the genome, the production of the target protein exceeded the levels achieved with the double plasmids, as indicated in Fig. 3.

**Fig. 3.**
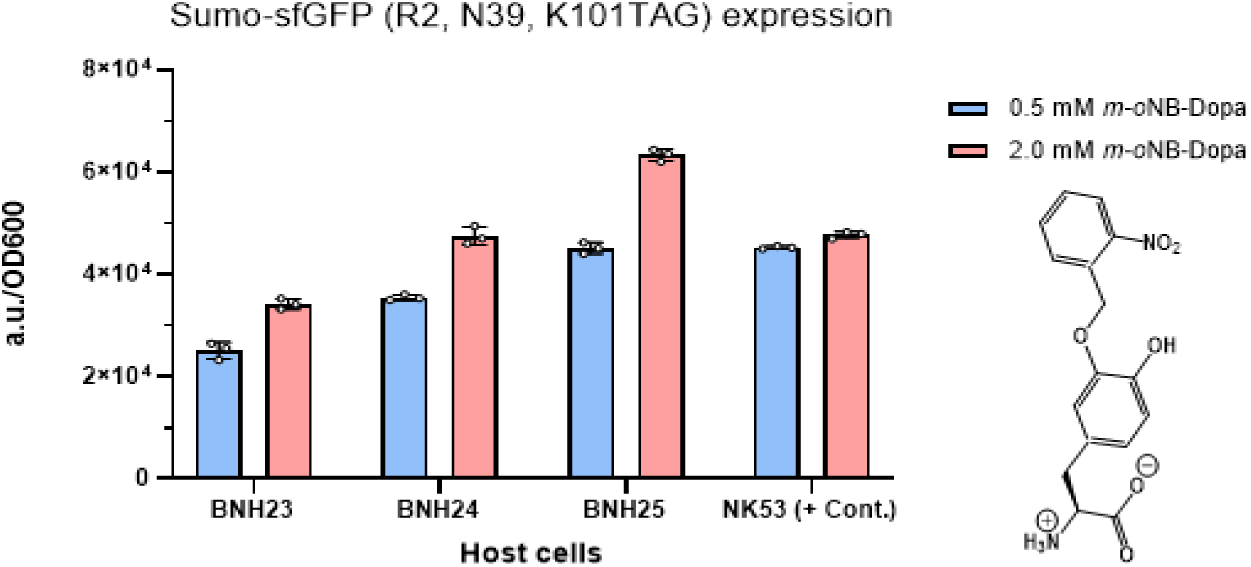
Calibrated fluorescence signal intensities, represented as fluorescence arbitrary units per OD_600_, for Sumo-sfGFP (3xAm) incorporating *m*-oNB-Dopa in the enriched strains. Each strain was cultivated in triplicate in 100 μL of ZYP-5052 medium with either 0.5 mM or 2.0 mM *m*-oNB-Dopa analog, and the cultures were incubated in a 96-well microplate overnight at 37°C and 300 rpm. The microplate was excited at 488 nm, and the fluorescence signal was detected at 530 nm using the Tecan Infinite 200 PRO plate reader. Bacterial growth was assessed by measuring absorbance at 600 nm. Right: Structure of *m*-oNB-Dopa (*meta*-(ortho-(2-nitrobenzyl))-3,4-dihydroxyphenylalanine). Error bars smaller than symbol width not shown. Data were presented as mean ± SD.

To further assess the efficiency of these strains, we examined the expression of the hybrid scaffold comprising an elastin-like protein (ELP) fused with sfGFP (ELP-sfGFP). This scaffold featured five, ten, twenty, and thirty Amber (TAG) codons. Among these strains, BNH25, with five integrated OTS cassettes, demonstrated remarkable efficiency in incorporating *m*-oNB-Dopa, yielding a production of over 100 mg/L of ELP(5xAm)-sfGFP. Moreover, proteins with 10 and 20 amber codons were also expressed at appropriate levels (Fig. 4). This underscores the reliability and robustness of BNH25 as a cell factory for the specific incorporation of *m*-oNB-Dopa into the desired proteins. We confirmed the full incorporation of the *m*-oNB-Dopa analogue through MS analysis of the purified ELP-sfGFPs featuring 5 and 10 amber codons (see Fig. S3).

**Fig. 4.**
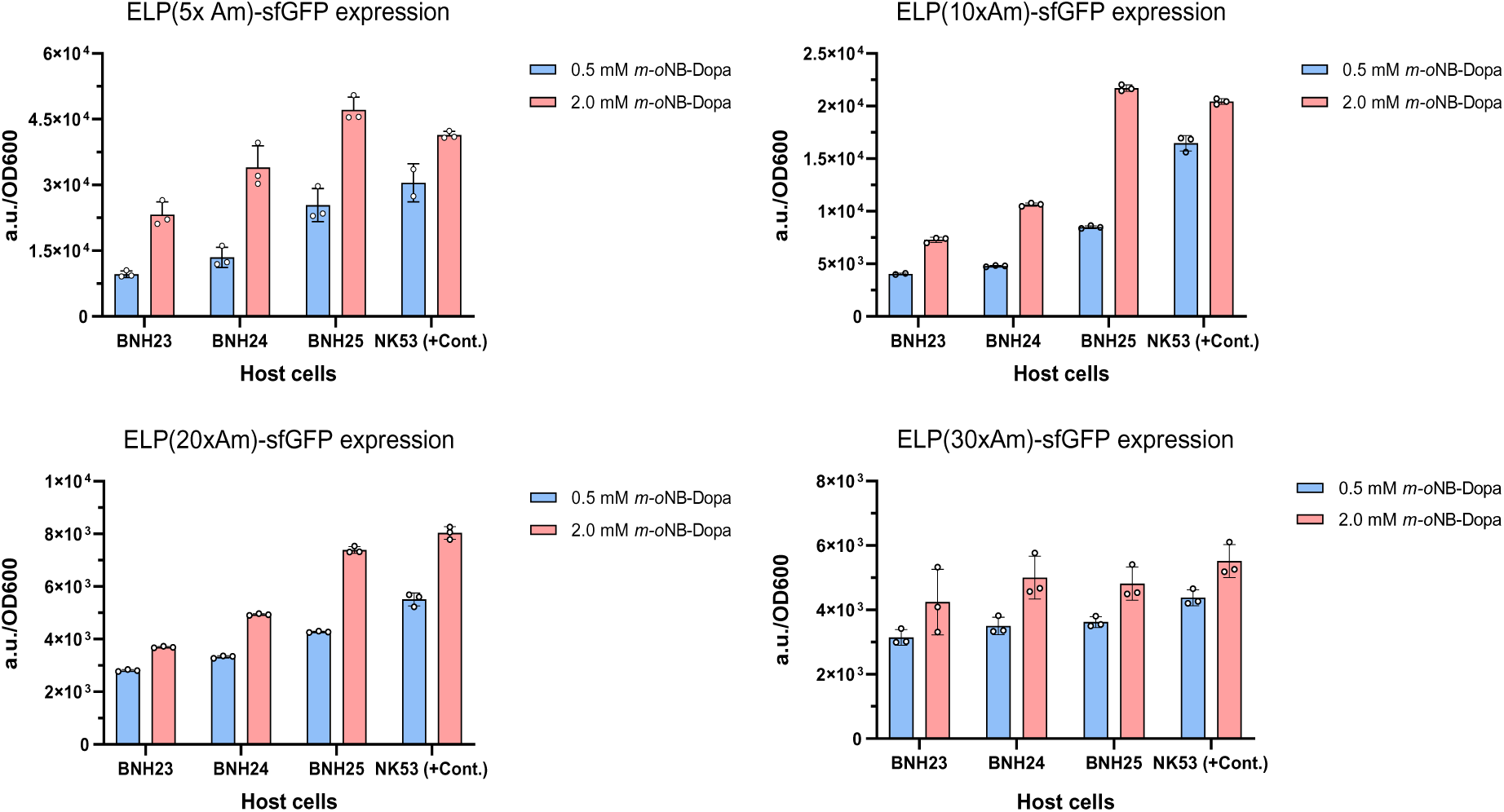
Analysis of ELP (n × TAG)-sfGFP (n = 5, 10, 20, and 30) production via fluorescence signal detection at 530 nm. Samples were excited at 488 nm using the Tecan Infinite 200 PRO. We employed 96-well plates to cultivate strains containing four distinct constructs, each featuring various Amber codons within the designed ELP. Each strain was cultured in triplicate using 100 μL of ZYP-5052 medium containing either 0.5 mM or 2.0 mM *m*-oNB-Dopa analogue. After an incubation period at 37°C and 300 rpm, we measured cell growth at 600 nm, and the calibrated signals were expressed as fluorescence arbitrary units per OD_600_. Error bars smaller than symbol width not shown. Data were presented as mean ± SD.

### Engineering an auxotrophic cell chassis with an integrated OTS to boost protein expression through stop-codon readthrough, utilizing fluoroproline analogs

To assess the impact of fluoroproline analogs on protein expression, we combined OTS with SPI. To create a cell chassis capable of accommodating both OTS and SPI, we introduced a *proB* gene null mutation in the BNH24 strain, resulting in the development of a new strain referred to MS34. This *proB* gene null mutation in BNH24 cells involved the insertion of a double stop codon and the deletion of eight amino acids within the glutamate 5-kinase gene sequence, leading to the generation of a stable proline-auxotrophic integrated OTS cell.

After confirming auxotrophy behaviour by growing the MS34 strain in minimal medium lacking proline, we evaluated the expression of ELP-sfGFP constructs in autoinduction minimal media. These media contained 2 mM *m*-oNB-Dopa along with 1 mM L-Pro (used as a positive control) or 1 mM of the 4*R*-FPro, 4*S*-FPro, and di-FPro analogs in 96-well plates. Our investigation focused on the influence of fluoroproline analogs during the translation of ELP (*n* × TAG)-sfGFP (*n* = 5, 10, 20, and 30), which contained 20% proline in the ELP segment (one in each pentapeptide, VPGxG motif) of the fusion construct. Expectedly, the incorporation of 4*R*-FPro analogs resulted in an increased production of ELP-sfGFP, even surpassing the levels achieved with L-Pro (refer to Fig. 5). Notably, the deletion of the *proB* gene in the BNH24 strain led to a decrease in ELP-sfGFP platform production compared to a ZYP-5052-rich medium, even in the presence of the canonical amino acid L-Pro (see Fig. S4). When we compared the ELP-sfGFP production capabilities of strain BNH24, we observed that the overexpression of the protein in minimal medium (NMM) decreased to less than 25%. However, the ELP-sfGFP production for both MS34 and BNH24 strains in the ZYP-5052-rich medium remained relatively similar (see Fig. S5). Interestingly, when we tested the impact of two analogs (*m*-oNB-Dopa and 4R-FPro), a positive effect of the 4R-FPro analogue on the translation of different ELP-sfGFP constructs was evident. This increased production of the target proteins was even observed in the non-auxotrophic strain BNH24 (refer to Fig. S6).

**Fig. 5.**
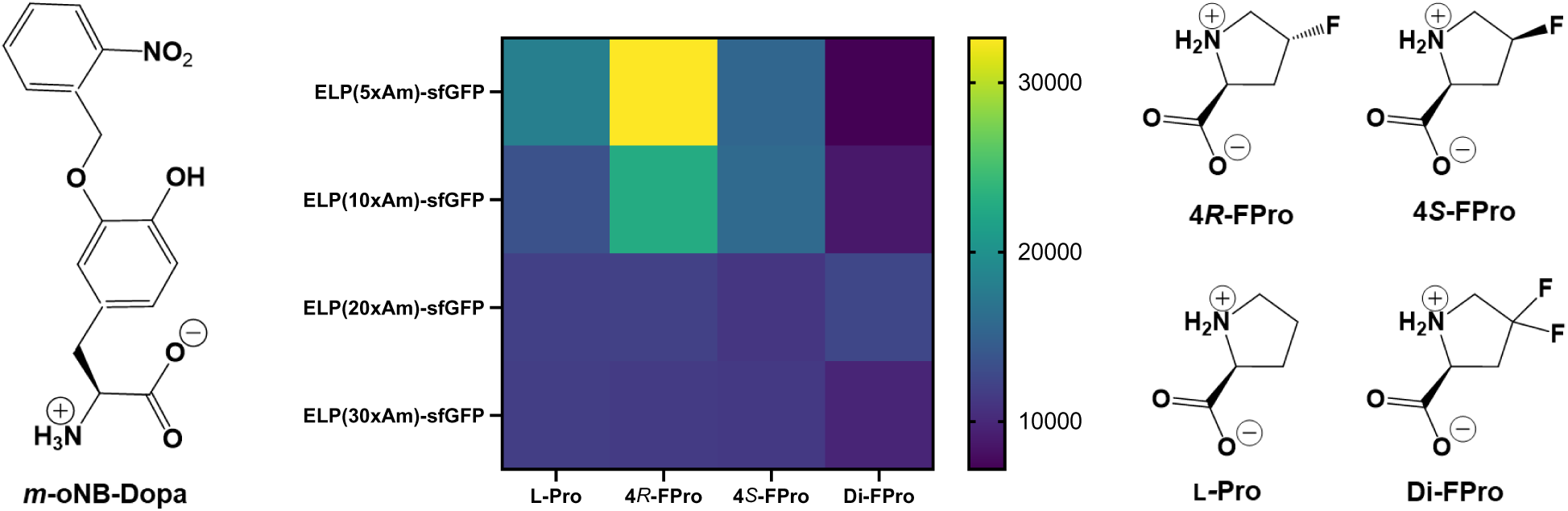
The OTS with auxotrophic protein expression: Effects of proline analogues on recombinant protein expression with stop codon readthrough. A heat map plot displays the expression levels of ELP-sfGFPs by the MS34 strain (a derivative of BNH24 with a *proB* gene deletion) in NMM autoinduction medium. The medium contains 1 mM L-Pro or various fluoroproline analogs, along with 2 mM *m*-oNB-Dopa. Proline (Pro) analogues are 4*R*-FPro: (2*S*,4*R*)-4-fluoroproline; 4*S*-FPro: (2*S*,4*S*)-4-fluoroproline; Di-FPro: (2*S*)-4,4-difluoroproline).

### Synthetic *m*-oNB-Dopa auxotrophism: establishment of a biocontained OTS-integrated cell chassis

To emphasize the superiority of the integrated OTS chassis over plasmid-based systems, we investigated the capability to create a biocontainated cell by introducing amber codons at specific sites within well-known essential genes in the *E. coli* genome. Efficient stop codon suppression either by natural suppressors or by o-pair are the ways to achieve functional rescue of these genes. Specifically, we targeted *murG* F213, *serS* F243, and performed deletions in the tyrT and tyrV tRNA genes, following the method by Rovner et al.^4^. This led to the generation of three strains: BNH24-*murG*::F213Am (MS36), BNH24-*murG*::F213Am, *serS*::F243Am (MS47), and BNH24-*murG*::F213Am, ΔTyr_VT_ (MS48). We cultured these cells in LB medium with and without *m*-oNB-Dopa in 96-well microplates. Analysis of the growth curves confirmed that the synthetic auxotrophic strains maintained robust growth in the presence of the *m*-oNB-Dopa analogue, even at low concentrations (62.5 μM).

To evaluate the specificity of the integrated OTS, we cultured the strains with L-Dopa, a naturally occurring compound sharing chemical properties. This compound results from UV-induced *m*-oNB-Dopa protecting group cleavage and serves as an isosteric analogue of canonical L-Tyr. The lack of growth over a 24-hour period at 32°C (as shown in Fig. 6) and the robust growth in the presence of 62.5 μM *m*-oNB-Dopa provide strong evidence supporting the biocontainment of strains MS47 and MS48.

**Fig. 6.**
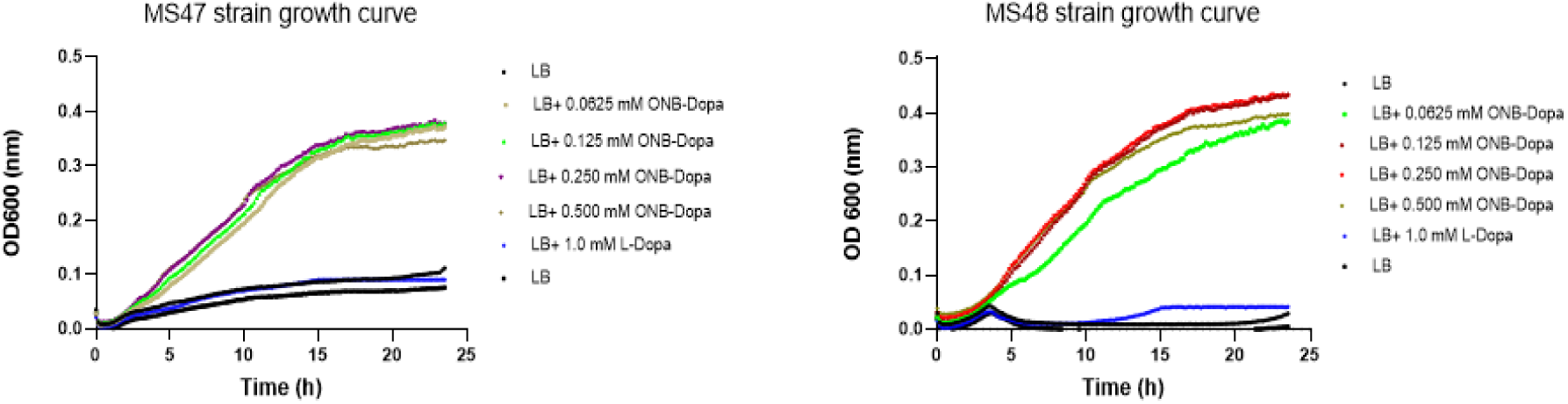
Adding biocontainment to the OTS integrated *E. coli* cells. Growth curve analysis of strains MS47 (BNH24-*murG* F213Am-*serS* F243Am) and MS48 (BNH24-*murG* F213Am-ΔTyr _VT_) in 96-well microplate at 32 °C. The Tecan Infinite 200 PRO was configured to measure absorbance every 10 minutes intervals following 2 minutes of shaking. To examine the effect, decreasing concentrations of the *m*-oNB-Dopa analogue were added fourfold to 100 μL LB medium. In addition, the orthogonality of the integrated OTS cassette with the _L_-Dopa amino acid was tested.

We also conducted experiments to measure the potential escape frequency of these strains by incubating the cells in LB medium at 37°C and 250 rpm for an extended duration. Among three separate cultures, only one batch of MS47 and MS48 exhibited weak turbidity after one week, with an OD_600_ of 0.3, compared to 5.0 in the positive control. The remaining cultures were further incubated and showed no signs of growth even after one month. Inoculating 10^9^ cells of the MS47 strain on LB agar plates in triplicate for one week resulted in an outbreak frequency of less than 10^-8^.

## Discussion

To realise the vision of a novel synthetic cell^27^, a durable genetic content is essential. The use of plasmid-based orthogonal translation systems to expand the genetic code has revealed significant limitations that hinder the wider application of this technology in various biotechnological contexts. To create robust biological networks, a variety of genetic circuits encompassing all essential components for recoded organisms is required. However, the extrachromosomal maintenance of genetic content poses major challenges for cells, such as vector instability, metabolic burdens, and replication control conflicts. To achieve optimal genetic fitness in a synthetic cell tailored to specific conditions, a stable genomic content is required that also allows for further robustness optimisation, e.g., by ALE experiments^28^. The transition from extrachromosomal genes to genome-based metabolic engineering therefore represents a promising opportunity to develop suitable platforms for the evolution of synthetic cells and to transition the orthogonal translation from academic research to practical industrial application.

Doubtless, by harnessing tools from the arsenal of bacterial-phage warfare, we are on the verge of modularizing bacterial or even eukaryotic genomes in a precise manner, paving the way for the creation of synthetic cells with tailored properties. We illustrate this here by successful generation of multicopy OTS integration into *E. coli* genome in a programmable and targeted manner, resulting in the development of a genome-recoded biocontained *E. coli* cell chassis. Conventional methods for genomic integration often rely on Cas9 double-strand break (DSB) activity coupled with λ Red recombination. However, this approach poses challenges due to its strong toxicity and lack of multiplicity. In contrast, the bacterial CRISPR-associated transposase tool (CASTs) offers a promising alternative. It allows for the insertion of large DNA sequences in a multiplexed manner without generating DSBs, mitigating toxicity concerns for cells.

Molecular machines facilitating the incorporation of non-canonical amino acids (ncAAs) are typically chosen by creating gene libraries of selected aminoacyl-tRNA synthetases. Two natural archaeal systems, the tyrosine pair *Mj*TyrRS:tRNA^Tyr^ from *Methanocaldococcus jannaschii* and the pyrrolysine pair from *Methanosarcina barkeri* or *mazei* species (*Mb/m*PylRS:tRNA^Pyl^), are tools of choice for the construction and selecting of suitable OTSs in *E. coli*. Orthogonal pairs (o-pairs) based on both the *Mj*TyrRS and PylRS scaffolds present advantages and disadvantages depending on experimental contexts. In bacterial host cells, *Mj*TyrRS often demonstrates superior performance compared to PylRS-based systems, such as yielding a higher amount of purified ncAA-modified target protein. PylRS-derived o-pairs in orthogonal translation typically yield a lower amount of target protein due to the enzyme’s generally low catalytic efficiency^29^. While o-pairs based on *Mj*TyrRS are usually efficient enough to repress a higher number of stop codons in frame, the same is not observed for orthogonal translation based on PylRS^30^. Consequently, the chosen OTS for the multiple incorporation of *m*-oNB-Dopa into our ELP scaffold was based on *Mj*TyrRS:tRNA^Tyr^ o-pair.

Using a dual plasmid system with a low copy number donor DNA plasmid and a moderate copy number pEffector plasmid under the control of the pTetA/R promoter, the entire CRISPR-associated transposon machinery from the type I-F CAST system derived from *Vibrio cholerae* Tn6677, we introduced an evolved *Mj*TyrRS:tRNA^Tyr^ o-pair [oNB-DopaRS:tRNA_CUA_]^31^ into the chromosome of an *E. coli* (DE3) B95 in which 95 of the 321 TAG[amber] codons were replaced by TAA[ochre] codons. Having eliminated the UAG-recognising termination factor (RF-1) along with five intracellular nitroreductases, the NK53 strain was equipped with a fivefold integrated OTS to decode the TAG amber codons for the synthetic amino acid *m*-oNB-Dopa. As a feature of NK53 strain, the deletion of the nitroreductase genes^20^ in the engineered cell suppresses the reduction of the NO_2_ group of *m*-oNB-Dopa analogues to ensure stability of oNB protective scaffold (prior to deprotecting). It is well known that the use of *m*-oNB-Dopa in mussel foot proteins to produce photoactive wet-bioadhesive materials as demonstrated elsewhere^31^. Controlled number of photoactivatable *m*-oNB-Dopa ncAAs in ELPs offer the possibility of producing a smart photo-controllable adhesive that is activated upon UV irradiation. The efficiency of *m*-oNB-Dopa incorporation in the BNH25 strain indicates that the CAST tool is capable of elevating gene expression at the copy-number level in a synthetic bacterial chassis. This chassis robustly grows and can overexpress protein congeners with coding sequences containing up to 30 in-frame stop codons^32^.

To enhance OTS systems, selective pressure incorporation (SPI) can be employed, involving the global replacement of canonical amino acids with synthetic analogs in auxotrophic cells^33^. This approach effectively addresses common issues associated with OTS, like the decline in translation efficiency with an escalating number of in-frame stop codons^34^. These analogs have the potential to impart novel spectroscopic and conformational properties to proteins, enhance protein stability and enzymatic activity, and even boost protein yield in comparison to the parent protein^35^. Therefore, considering the strengths and weaknesses of both the SPI- and OTS-based methods, there’s a strong motivation to merge them in a single *in vivo* expression experiment^33^. This combination offers the advantage of using OTS to introduce ncAAs at specific, permissive sites while using SPI to incorporate structurally similar ncAAs in response to sense codons with multiple residues (i.e., to substitute canonical amino acids with non-canonical ones) generating a simple protein labelling (see scheme 1). Genome integration is a powerful tool to robustly merge SPI-mediated substitutions of proline residues in ELP-Pentamers with exocyclically modified analogues to manipulate the protein backbone^36^ with OTS for site-specific incorporation of the aromatic ncAA *m*-oNB-Dopa, which is known to be a crucial component for photo-controlled wet bioadhesion properties^37^. As can be see in Figure 5 the OTS with auxotrophic protein expression is significantly enhanced, as mirrored in the corresponding protein yields.

Given that the ELP constructs contain a pentapeptide polymer with one Pro-residue/pro monomer in the second position (20% of the sequence), we anticipated that manipulating the pyrrolidine ring’s physicochemical properties would impact protein expression and yield. For instance, replacing all prolines in the ELP pentamer with the 4*R*-FPro analogue using the SPI method (in minimal medium) was expected to increase the ribosomal translation rate. Conversely, the 4*S*-FPro analogue was predicted to have a negative effect on protein expression. Through the generation of Pro-auxotrophic OTS-integrated cells (BNH24, Δ*proB*), an analysis of ELP-sfGFP expression was conducted in minimal media containing various fluoroproline analogues. While the expression level decreased in the BNH24 (Δ*proB*) strain in minimal media, the positive effect of the 4*R*-FPro analogue on the expression of ELP (5x and 10x stop codons)-sfGFP was significant (more than 80%). In contrast, the negative impact of the 4*S*-FPro analogue was evident, even in non-auxotrophic, integrated OTS cells (see Fig. S6).

To ensure safe bioproduction of such high-value materials, we used the CRISPR-assisted genome editing tool to insert TAG codons instead of two Phe sense codons into essential genes of our chassis to create an orthogonal barrier between the engineered cells and the environment (genetic isolation). Although the engineered *E. coli* strains (MS47 and MS48) exhibit strong biocontainment behavior, with undetectable escape frequencies during culturing in lysogeny broth medium, the reinforcement of such safeguards can be easily achieved by increasing the number of TAG codon mutations in conserved functional residues of essential genes. The specificity of the engineered enzyme [oNB-DopaRS:tRNA_CUA_], highlighted by the absence of bacterial growth in the presence of L-Dopa, the most isosteric amino acid to *m*-oNB-Dopa, underscores the robustness of synthetic *m*-oNB-Dopa auxotrophism in our cell chassis. Assessing bacterial growth in the presence of lower synthetic amino acid concentrations highlights the engineered cells’ robustness. This positions them as a viable bacterial cell factory for producing proteins with *m*-oNB-Dopa incorporation, suitable for demanding applications in biomaterial research. Robust and safe chassis with specific OTSs are essential to create functional recoded organisms suitable as biotechnological production platforms.

The efficient integration of our orthogonal translation systems into the *E. coli* genome supports the hypothesis of generating synthetic bacterial cells with a recorded genome that grow under controllable conditions and are of great biotechnological value as potential production platforms with built-in biosafety^28^. Anticipating the expansion of the genetic code using such platforms, we foresee the cost-effective production of new-to-nature peptides and proteins through the strategic integration of different OTS modules (o-pairs with compartmentalization signals, engineered elongation factors, orthogonal ribosomes with correct rRNA modifications etc.). This also involves targeting additional rare codons of the microbial genetic code such as AGT, TCA, TCG (Ser codons), CTA (Leu codon), or AGA and AGG (Arg codons), thereby establishing whole cells or compartmentalised modular biosynthetic gene islands with altered/alienated genetic codes.

However, the significant biotechnological, medicinal, and pharmacological applications of OTS can be fully realized only when accompanied by synthetic metabolism – the in-situ biosynthesis of synthetic biochemical building blocks or analogues. Currently, the absence of such platforms stands as the primary impediment to the economic large-scale production of synthetic peptides and proteins. In this context, our cellular chassis, featuring stable and robust genomic integration of OTS along simple metabolic markers and built-in biosafety, stands as a promising prototype for future microbial cell factory platforms. These platforms hold significant appeal for the production of valuable smart biomaterials, therapeutic proteins, and other high value bioproducts. They operate within a genetically isolated environment with inherent biocontainment measures. Therefore, we envision our genomically integrated, robust, modular and biosafe recoded chassis as promising microbial cell factories of the future.

## Methods

### Plasmid construction

All plasmids and primers used in this study are listed in Supplementary Tables S1-3. The pTet effector array1 plasmid construct (PM452) was cloned by generating PCR fragments from the pTetQCas-8+IS186 and the pTnsABC construct (gift from Sheng Yang (Addgene plasmid # 170636; & #1306331)) using the GeneArt™ Gibson Assembly HiFi Master Mix (InvitrogenTM). pDo-ColA-RE-TacI-oNB-DopaRS-tRNA_CUA_-LE donor plasmid was prepared in two steps. First, pDonor-oNB-DopaRS was cloned by amplification of vector backbone including mini-transposon ends from pDonor-GDH (gift from Sheng Yang (Addgene plasmid # 140629)) and generating a PCR fragment of oNB-DopaRS gene cassette (containing pTacI-oNB-DopaRS+ ProK promoter-tRNA_CUA_ optimized sequence) from pUltra-oNB-DopaRS (homemade construct) using Gibson assembly approach. Then, pDo-ColA-oNB-DopaRS-tRNA_CUA_ (PM453) plasmid was cloned using PCR amplified fragments from the pTnsABC plasmid backbone and oNB-DopaRS mini-Tn fragment (Fig. 1). For site-specific genome editing or gene disruption for auxotrophic cell creation, the pEcCas and pEcgRNA^17^ (was a gift from Sheng Yang (Addgene plasmid # 73227; & #166581)) were applied while the NEB Golden gate cloning kit was used to build appropriate pEcgRNA-N20 plasmids. All PCR fragments for cloning were generated using Phusion™ High-Fidelity DNA Polymerase (Thermo Scientific^TM^). All primers and oligonucleotides as donor DNA were synthesized by Integrated DNA Technologies (IDT) Company.

To construct pEcgRNA-N20 plasmids with different spacers, the original pEcgRNA (Addgene#166581) containing the *ccdB* gene with BsaI sites at the ends was digested with the restriction enzyme BsaI to generate linearised pEcgRNA with the 5′-TAGT-3′ and 5′-AAAC-3′ overhangs; the linearised pEcgRNA was either used immediately or frozen at −20 °C for later use. The synthesized single strand oligonucleotides (24 bp) comprising of 4 nt of the overhangs and 20 nt of target specific sequences (spacers) were annealed to form dsDNA, which was then ligated to BsaI-linearized pEcgRNA to generate the new target specific plasmid. A reaction mix consisting of 14 μl ddH_2_O, 2 μl T4 ligase buffer (10x), 2 μl (20 μM) N20-up, and 2 μl (20 μM) N20-dn was incubated at 95 °C for 5 min, and the temperature was then gradually reduced in a thermocycler (Mastercycler® nexus gradient; Eppendorf) and held at 16 °C for 10 min. The annealed dsDNA was then diluted 200-fold and 1 μl of the diluted DNA was ligated to 50 ng of BsaI-linearized pEcgRNA in a 20 μL mixture of T4 ligase buffer, and T4 ligase for 30 min at 25 °C. The ligation product is transformed into DH5α chemically competent cells to select colonies harboring plasmid pEcgRNA-N20. Positive clones were selected on LB plates supplemented with 50 μg/ml spectinomycin and plasmids were extracted after overnight culture of two or three isolates, using GeneJET Plasmid Miniprep Kit (Thermo Scientific^TM^). Authenticity of the constructs were confirmed by Sanger DNA sequencing (GENEWIZ, Azenta Life Sciences).

### *E. coli* culturing, transposition and genome editing experiments

To achieve integration of an oNB-Dopa orthogonal translation system into a suitable engineered host cell, a modified *Escherichia coli* strain, NK53, was used. *E. coli* strain B-95.ΔAΔfabR^18^ (was provided by the RIKEN BRC through the National BioResource Project of the MEXT, Japan (cat. RDB13711)) with no specific assignment to the UAG codon was previously modified by deletion of six nitroreductase genes to generate an attenuated nitroreductase activity strain (NK53) responsible for reduction of the nitro group on *m-o*NB-Dopa incorporated expressed protein. The genotype of new modified *E. coli* strain is as follow: *E. coli* B-95.ΔAΔfabR; ΔNfsA::FRT; Δ*NfsB*::FRT; Δ*AzoR*::FRT; Δ*Ydja*::FRT; Δ*NemA*::FRT; Δ*RutE*::FRT, hereinafter is called NK53 strain^20^. A full list of *E. coli* strains used for genome editing experiments is provided in Supplementary Table S3.

All *E. coli* transformations were performed using homemade chemically or electro competent cells and standard heat shock transformation or electroporator, followed by recovery in LB at 37 °C and plating on LB agar media with the appropriate antibiotics at recommended concentrations. Transposition experiment was performed after confirming the authenticity of PM453 (pDo-ColA-RE-TacI-oNB-DopaRS-tRNA_CUA_-LE) plasmid. PM452 and PM453 plasmid constructs were co-transformed into NK53 strain using chemically competent cells. After 90 min recovery time in LB or SOB medium at 37 °C, total cells were spread on LB+Kan.+Spect. plate and incubated overnight. The grown colonies were scraped from the plate and resuspended in sterile ddH_2_O; then a suitably diluted cell sample was spread on LB+Kan.+Spect. applying 0.5 ng/ml anhydrotetracycline (aTc) as transposition inducer. Second round of transposition was conducted by spread a resuspended scraped cells on LB+Kan.+Spect.+5.0 ng/mL aTc to increase number of integrated OTS (Mini-Tn: oNB-DopaRS gene cassette) in NK53 cells. Three isolates were picked up from each plate and cultured in LB+Kan liquid media overnight to make glycerol stock. Colony PCR was performed in parallel to detect insertion of the OTS using 9 pairs of primers which are flanking all nine target sites (designed in crRNA array) in NK53 *E. coli* genome (Supplementary Table S3). Proper size of PCR product confirms the integration of the target OTS after agarose gel visualization.

For genome editing (site-specific mutation, or gene disruption) experiments, after cloning of appropriate pEcgRNA-N20 plasmid, the target *E. coli* strain is transformed by the pEcCas plasmid (PM272) which contains Cas9 and λ Red system genes and plated on LB+Kan. A single colony is cultured overnight; then inoculated into a new LB+Kan liquid medium to prepare electrocompetent cells. 30 min before reaching growth to OD 0.3-0.4, 10 mM Arabinose is added to induce λ Red enzyme expression. 100 ng corresponding pEcgRNA-N20 and 400 ng donor DNA were co-electrotransformed into 100 uL competent cells in 1 mm gap electroporation cuvette by applying 2.5 kV pulse in electroporator (Eppendorf™ Eporator™). After adding 900 uL of cold SOC medium, the cells were recovered and spread on LB+Kan.+Spect. plate overnight. Three colonies were randomly chosen and cultured overnight in LB+Kan. + 10 mM L-Rhamnose. One can keep these strains in glycerol stock format if one wants to continue genome modification by targeting other genes. L-Rhamnose induces a gRNA transcription under control of rhaBAD operon in pEcCas to target ColE1 ori in pEcgRNA plasmid and eliminate it.

Colony PCR was conducted to amplify target gene regions using designed primer pairs flanking the target site (~200 bp upstream and downstream). PCR products were visualized by 1% agarose gel, purified, and sent for Sanger DNA sequencing. Similar PCR reaction was done for ancestor strain to compare and confirm the genome editing.

### Plasmid curing

For plasmid elimination in OTS integrated strains, the pCutamp plasmid (gift from Sheng Yang (Addgene plasmid # 140632)^38^) containing a gRNA sequence under the rhamnose-inducing promoter system that targeting the AmpR promoter in the PM452 and PM453 plasmids was chemically transformed. After 1 hr recovery at 37 °C, 0.3 mL recovered cell culture was reached to 0.9 mL LB+50 μg/mL Apr.+10 mM L-Rhamnose culture and incubated at 37 °C and 250 rpm for 5 hours. 100 μL cell sample (10^4^-times dilution of the culture) was spread on LB+Apr.+ 5 mM L-Rhamnose plate and incubated overnight. Single colony picked up for each strain and cultured in 0.5 mL LB+10 mM sucrose liquid medium overnight. Then, 100 μL cell samples (10^6^-times diluted culture) were plated on LB+Sucrose to get a single colony. pCutamp carries the *SacB* gene, the product of which can convert sucrose to levan, which is highly toxic to *E. coli*. The plasmid elimination in cured strain was tested by spotting individual colonies (3-5 colonies) on LB+Kan., LB+Spect., and LB+Apr., and LB plates, simultaneously (Fig. S3). Strains grown on the LB plate but not on other antibiotic-containing plates revealed proper plasmid curing. The cured strains are grown in LB media to make glycerol stock or competent cells.

For plasmid curing in CRISPR-Cas9-assisted genome edited strains using pEcCas (PM272) and pEcgRNA-N20 plasmids, after confirming authenticity of genome editing by Sanger DNA sequencing, glycerol stock of the edited strains is cultured in LB+Kan+10 mM L-Rhamnose overnight. 100 μl cell samples (10^6^-times dilution of the culture) were spread on LB+Kan.+5 mM L-Rhamnose and incubated overnight. The same strategy was designed for targeting the AmpR promoter in pEcgRNA-N20 plasmids by induction of specific gRNA in pEcCas plasmid. Single colony was picked up and cultured in LB+ 10 mM Sucrose liquid medium for 6 hours at 37 °C. 100 μl cell samples (10^6^-times dilution of the culture) were plated on LB+sucrose and incubated overnight. The colonies grown on plate were tested for kanamycin and streptomycin sensitivity to confirm proper plasmids elimination. pEcCas carries the *SacB* gene, and it will be eliminated in the presence of sucrose. Regarding TAG amber codon substitution in *murG* F243 and s*erS* F213 positions, all liquid and solid cultures were supplemented by 0.5 mM *m-o*NB-Dopa analog to support biocontained strain growth.

### Incorporation efficiency analysis

96 well microplate culture was performed for *m*-oNB-Dopa incorporation efficiency assessment of the integrated OTS chassis. NK53 strains integrated with a series of *m-o*NB-Dopa o-pair cassette from one to five copies were transformed with the plasmids containing Sumo-sfGFP (1x amber), Sumo-sfGFP (3x amber), ELP(5xamber)-sfGFP, ELP(10xamber)-sfGFP, ELP(20xamber)-sfGFP, and ELP-(30xamber)-sfGFP in parallel with a plasmid-based overexpression of OTS and target model proteins in ancestor NK53 strain as positive control and grown in 100 uL ZYP-5052 autoinduction medium supplemented by 0.5, 1.0. and 2.0 mM *m-o*NB-Dopa analog at 37 °C, 300 rpm for 24 hours. The incorporation efficiency can be measured by fluorescence signal measurement of the expressed sfGFPs harboring various numbers of amber stop codons as a sign of efficient incorporation using Tecan plate reader (Infinite 200 PRO). The 10 mL ZYP-5052 medium consisting of 2 mM *m-o*NB-Dopa was used for production of higher amounts of the ELP (5 or 10xAm)-sfGFP. High-performance immobilized metal affinity chromatography (IMAC) column (HisTrap^TM^ HP, cytiva) was used to purify expressed proteins from lysate in buffer H1 (20 mM Tris, 200 mM NaCL, 0.02% NaN_3_, pH 8.0). The loaded sample was washed by 45 mM imidazole in H1 buffer and the His6-taged protein was eluted by 300 mM imidazole in H1 buffer.

### ESI-MS Analysis

The Ni-purified ELP-sfGFP variants were dialyzed against 5 mM ammonium bicarbonate, pH 7.8 and directly infused in Electrospray ionization time of flight mass spectrometry (ESI-TOF-MS) by autosampler with Bruker Elute system at flow rate 0.3 mL/min. The proteins were ionized in positive ion mode applying a cone voltage of 40 kV while scanning from 300 to 3000 m/z. Raw data were analyzed employing the maximum entropy deconvolution algorithm with Bruker Data Analysis.

### Generation of biocontainment strain

To do point mutation for replacing phenylalanine codon (TTT) with TAG amber codon at *murG* F243 and s*erS* F213 position, appropriate gRNA sequences were designed as well as 89 bp single strand oligonucleotides as donor DNA (see Table S2 for detail). Initially, strain BNH24 was genomically edited by substitution of a single TGA codon at *murG* F243 position to create strain MS36 (BNH24-*murG*::F243TAG). Then further TAG replacement was performed in the MS36 strain at *serS* F213 position or null mutation of tRNA Tyr_VT_ to increase dependency of the strain on synthetic amino acid, oNB-Dopa.

Cell survival and escape from liquid cultures was monitored by quantifying c.f.u. on permissive (LB+*m-o*NB-Dopa) and non-permissive (LB) solid media, respectively. To quantify the degree of containment, the ratio of colony-forming units (c.f.u.) on non-permissive to permissive solid media was measured and cell survival and escape frequencies were calculated.

### Next-Generation Whole Genome Sequencing

The MS47 strain (BNH24-*murG*.F243xAm; *serS*.F213xAm) was chosen for next-generation whole genome sequencing by SeqCoast Genomics LLC, after culturing of the strain in permissive medium and extraction of the genomic DNA by PureLink™ Microbiome DNA Purification Kit (ThermoFisher Scientific). Short read whole genome sequencing, Illumina, by 400 Mbp/2.7 million reads provides 150 bp paired end reads for analysis. Breseq as a computational pipeline for finding mutations relative to a reference sequence for haploid microbial-sized genomes was applied to find indels and point mutations using B95.deltaA-CP085842 and PM453 plasmid as references. Breseq reports single-nucleotide mutations, point insertions and deletions, large deletions, and new junctions such as those produced by trasnposons in an annotated HTML format^39,40^. We used the Digital Research Alliance of Canada server to run breseq. Summary of the analysis has presented in Fig. S7.

### Growth curve analysis

The OD600 growth curve analysis was performed by inoculating target cells or isolated auxotrophic strains from −80 °C glycerol stocks into LB media for overnight growth. Then, 1% of each culture was inoculated in a 96-well cell culture plate in the desired growth media. Growth assays were then performed with a Tecan Infinite 200 PRO plate reader shaking at 32 °C for 24 h, with an OD at 600 nm taken every 10 min. Each sample was measured in three technical replicates in separate wells on the sample plate, and values were normalized to blank wells containing media only.

### Statistical analysis

All fluorescence intensity and culture growth measurements were performed in three independent triplicate samples and the data were statistically analysed by GraphPad prism 8.0.2 software.

## Acknowledgments

This research was supported by Canada Research Chairs Program (Grant No. 950-231971). We are very grateful to Chun Hin Wong and Department of chemistry facility room for the support in ESI-MS measurements.

