## Supplementary figures for "Genomically integrated orthogonal translation in *Escherichia coli*, a new synthetic auxotrophic chassis with altered genetic code, genetic firewall, and enhanced protein expression"

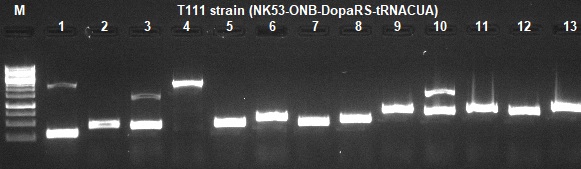

BNH21

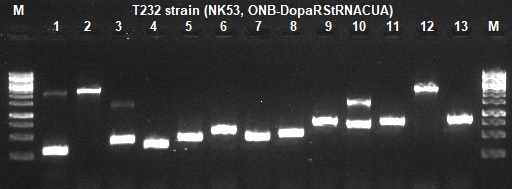

BNH22

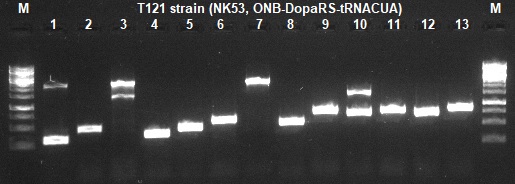

BNH22-2

BNH23

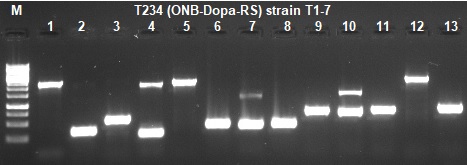

**Supplementary Figure 1**. **PCR analysis of the isolated strains after the transposition experiment using a pair of primers flanking the target genome regions.** BNH is the abbreviation for *E. coli* B strain and reductase-deleted cells. #2 stands for the integration of the oNB-DopaRS cassette. # The second digit stands for the number of OTS cassette insertions. It is worth noting that the presence of the additional PCR bands at sites #4, 7 and 10 (IS4-R2) indicates non-specific PCR products (site #4, 7, and 10 in BNH23). Wells #9-13 show PCR analysis of five similar target sites of the IS4-like element in the *E. coli* genome. Further experiments showed that only the fourth IS4 element is capable of integration (site #12). Strains BNH22 and BNH22-2 are two forms of double OTS-integrated cells. The size of the cargo is 2076 bp. PCR product sizes are the cargo size plus upstream and downstream regions depending on designed primers. DNA ladder is GeneRuler 1 kb, Thermo Fisher Scientific Inc., SM0313.

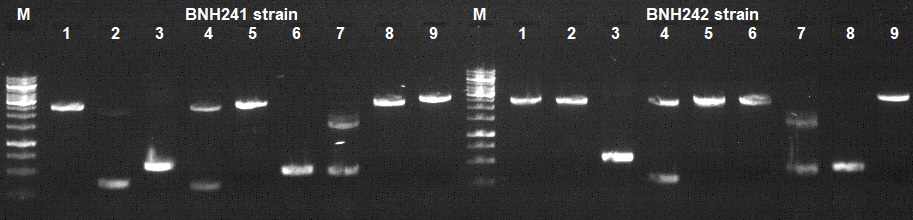

BNH24

BNH25

**Supplementary Figure 2**. **PCR screening of the new isolated strains after the re-transposition experiment in BNH23 strain.** Well #9 corresponds to the fourth site of the IS4-like element in the *E. coli* genome. The size of the cargo is 2076 bp. PCR product sizes are the cargo size plus upstream and downstream regions depending on designed primers. The DNA ladder is GeneRuler 1 kb, Thermo Fisher Scientific Inc., SM0313. Extra bands at sites 4 and 7 are non-specific products.

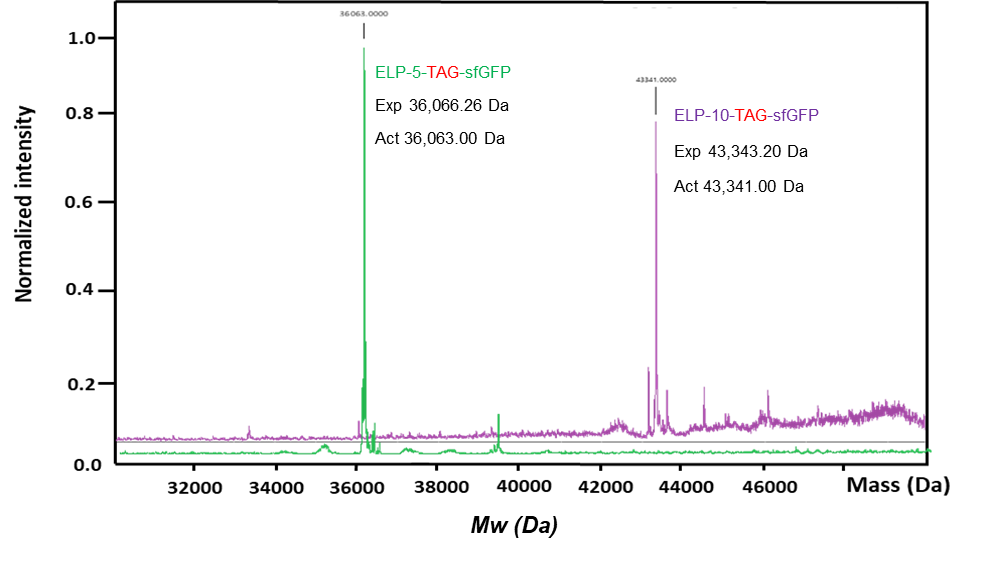

**Supplementary Figure 3.** ESI-MS analysis of the expressed ELP(5xAm)-sfGFP and ELP(10xAm)-sfGFP proteins after HisTrap HP-column purification. The molecular mass peaks of proteins confirm full incorporation of *m*-oNB-Dopa into the expressed ELP proteins.

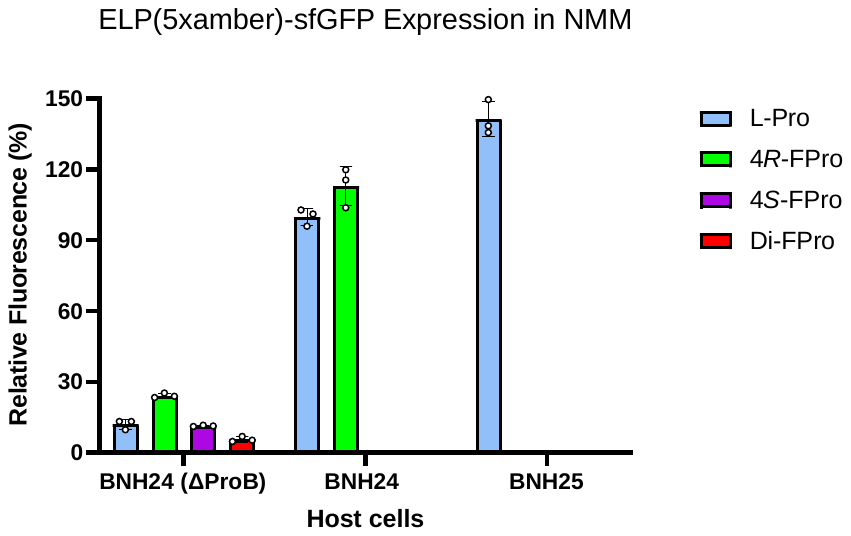

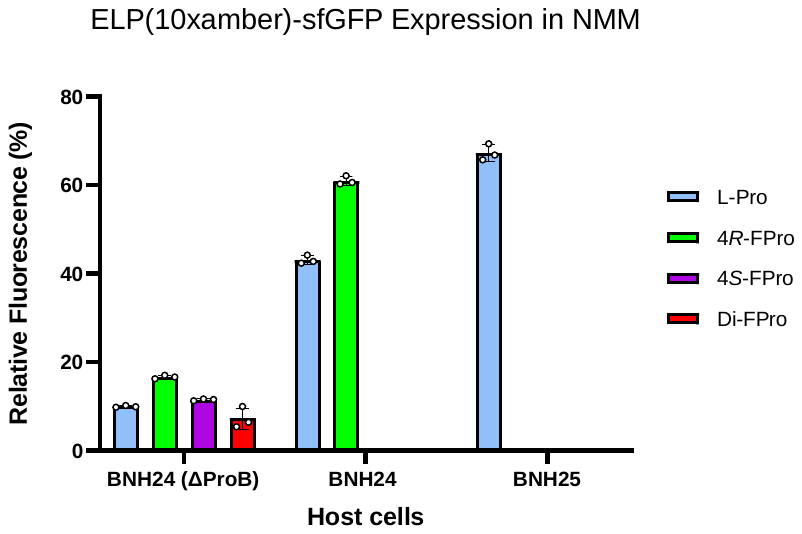

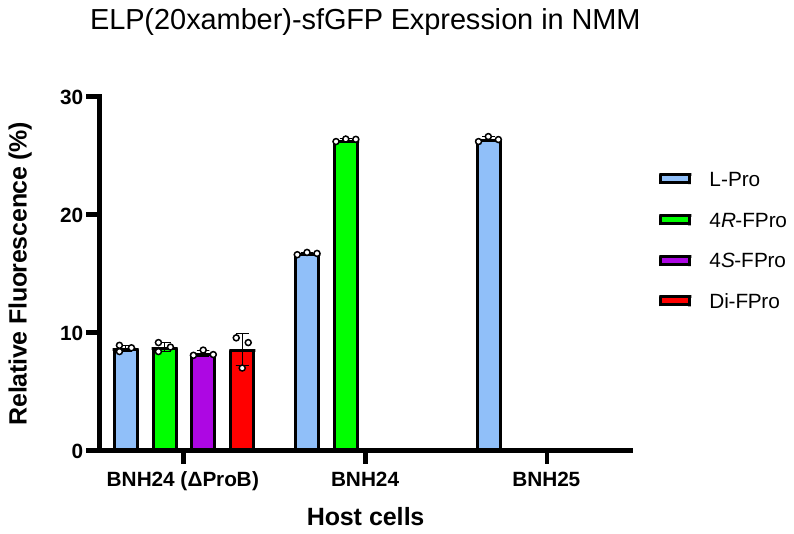

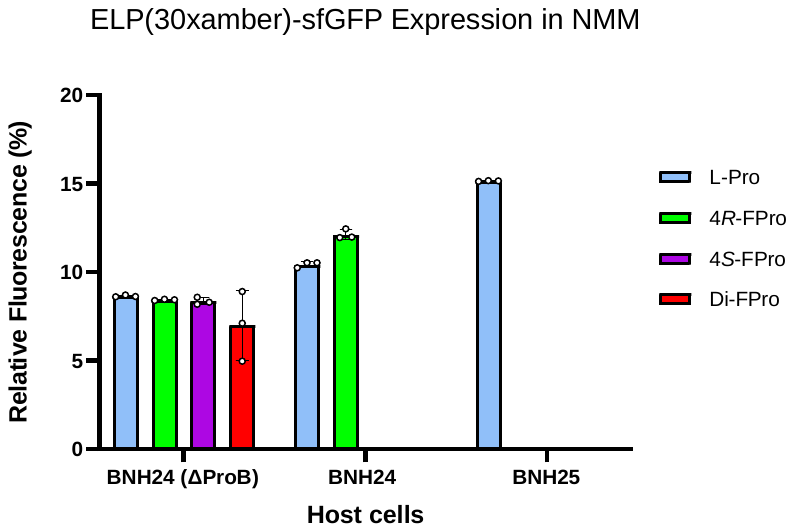

**Supplementary Figure 4**. **Combination of OTS and SPI to evaluate the effect of fluoroproline analogues**. Expression analysis of various ELP-sfGFP constructs in 100 μL NMM autoinduction media containing 1 mM fluoroproline analogs and 2 mM *m-o*NB-Dopa. The expression efficiency of the BNH24-Δ*proB* (MS34) strain with the ancestral non-auxotrophic BNH24 and BNH25 strains in the presence of the 4*R*-FPro analogue was compared. Proline (Pro) analogues are 4*R*-FPro: (2*S*,4*R*)-4-fluoroproline; 4*S*-FPro: (2S,4S)-4-fluoroproline; Di-FPro: (2S)-4,4-difluoroproline.

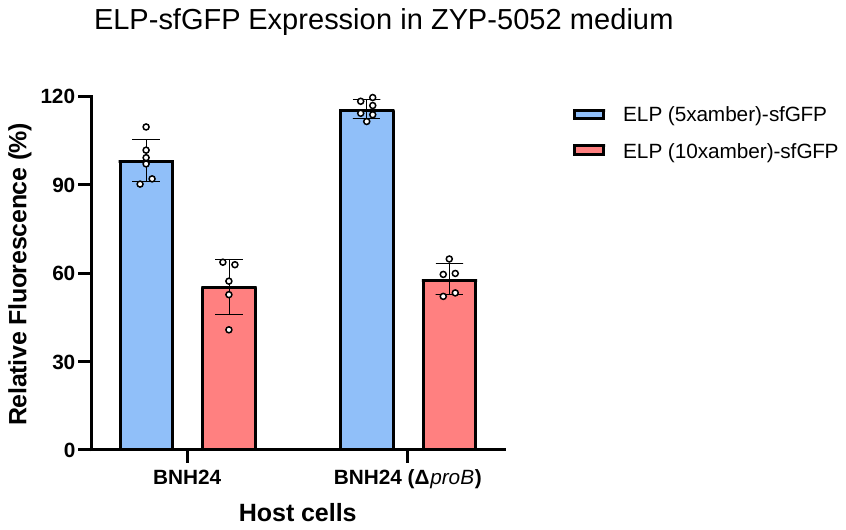

**Supplementary Figure 5**. **Analysing the effect of the null mutation of the *proB* gene in the genome of strain BNH24 on ELP (5 or 10xAm)-sfGFP expression in ZYP-5052 rich medium**. Comparison of the effects of the number of in-frame stop codons on the expression of recombinant proteins in the auxotrophic chassis with the original genomically integrated OTS strain in a rich medium.

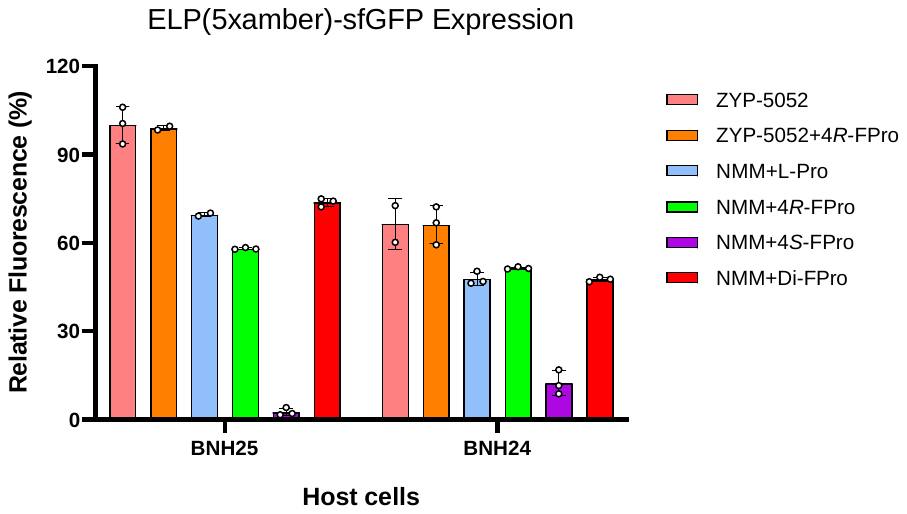

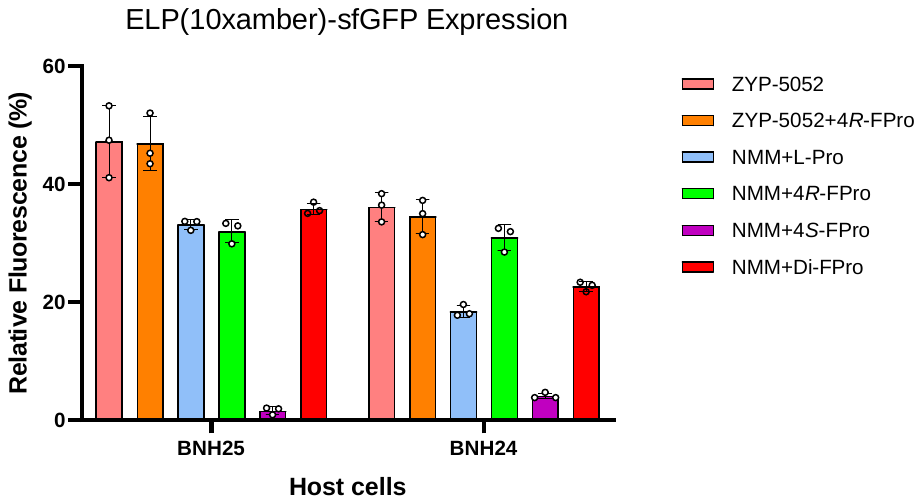

**Supplementary Figure 6**. **Positive effects of the 4*R*-FPro analogue on the performance of the OTS system**. Evaluation of the expression of ELP (5 or 10xAm)-sfGFP in both ZYP-5052 and new minimal media (NMM), incorporating various fluoroproline analogues. These assessments were conducted using non-auxotrophic strains, specifically BNH24 and BNH25. A positive impact of the 4*R*-FPro analogue on ELP-sfGFP production was observed in NMM, particularly in non-auxotrophic cells like BNH24. Expectedly, when fluoroproline analogs were introduced during the initial growth step upon inoculating non-auxotrophic BNH24 and BNH25 cells, the 4*S*-FPro analogue exhibited a strong inhibitory effect on protein expression (as it is known to be an inducer of ribosomal stalling^24^).
